# Systems genetics identifies modifiers of Alzheimer’s disease risk and resilience

**DOI:** 10.1101/225714

**Authors:** Sarah M. Neuner, Timothy J. Hohman, Ryan Richholt, David A. Bennett, Julie A. Schneider, Philip L. De Jager, Matthew J. Huentelman, Kristen M. S. O’Connell, Catherine C. Kaczorowski

## Abstract

Identifying genes that modify symptoms of Alzheimer’s disease (AD) will provide novel therapeutic strategies to prevent, cure or delay AD. To discover genetic modifiers of AD, we combined a mouse model of AD with a genetically diverse reference panel to generate F1 mice harboring identical ‘high-risk’ human AD mutations but which differ across the remainder of their genome. We first show that genetic variation profoundly modifies the impact of causal human AD mutations and validate this panel as an AD model by demonstrating a high degree of phenotypic, transcriptomic, and genetic overlap with human AD. Genetic mapping was used to identify candidate modifiers of cognitive deficits and amyloid pathology, and viral-mediated knockdown was used to functionally validate *Trpc3* as a modifier of AD. Overall, work here introduces a ‘humanized’ mouse population as an innovative and reproducible resource for the study of AD and identifies *Trpc3* as a novel therapeutic target.

**Highlights:**

- New transgenic mouse population enables mapping of AD risk and resilience factors
- Transcriptomic and phenotypic profiles in diverse AD mice parallel those in humans
- *Apoe* genotype and expression correlate with cognitive symptoms in mice
- *Trpc3* is a novel target to reduce amyloid load and cognitive symptoms in AD

## Introduction

Alzheimer’s disease (AD) is a neurodegenerative disorder characterized by both dementia and the accumulation of neuropathological amyloid plaques and tau tangles (Selkoe, 1991). Mutations that drive overproduction of beta-amyloid (Aβ) have been shown to cause early onset familial AD (FAD), leading to a model in which production and accumulation of Aβ is thought to be an initiating event in a sequence leading to memory loss, neurodegeneration, gliosis, and synaptic dysfunction (Hardy and Higgins, 1992). However, strategies to directly target amyloid for clearance have not translated into successful treatments and the number of deaths attributable to AD as well as costs associated with the disease continue to rise. In addition, even among patients with FAD mutations, the age at first symptom onset is widely variable, with some patients exhibiting symptoms decades later than predicted based on mutation status, suggesting additional genetic factors exist that may provide protection from disease (Ryman et al., 2014).

Although age is the greatest risk factor for developing AD, it is increasingly clear that genetics and family history play a large role. The heritability of AD is estimated to be in the range of 50-80%, indicating an individual’s susceptibility or resilience to disease is, at least in part, determined by genetic makeup (Gatz et al., 1997). Indeed, a number of genetic risk factors have been identified in recent years (Lambert et al., 2013). However, besides *APOE* and *TREM2*, effect sizes of common variants are generally small and a large proportion of the heritability remains unexplained by currently identified variants (Ridge et al., 2013). In addition, the study design utilized in most genome wide association tests (e.g. case-control) is primed to identify variants associated with *risk* of AD, rather than variants that modify an individual’s disease trajectory and/or delay the onset of disease and provide protection from AD. This is due primarily to the fact that asymptomatic individuals rarely enter the clinic for treatment, and even if they are included in a study, are likely to be enrolled as cognitively normal controls.

Identifying genetic variants and pathways involved in protection from AD will provide valuable targets for new therapeutics to prevent or delay the onset of symptoms. Individuals with high-risk genotypes but who fail to present with clinical symptoms of AD represent an ideal population in which to study resilience. However, causal mutations in *APP* and *PSEN1/2* (i.e. high-risk genotypes) are rare in humans, greatly limiting statistical power and opportunity for observation. In addition, access to brain tissue at early disease time points, before overt symptom onset, is limited for obvious reasons in human studies, precluding the identification of causal molecular mechanisms. Thus, mouse models harboring causal AD mutations are an important tool that present many advantages including defined high-risk genotypes, early access to brain tissue, and precise environmental control.

Recent arguments concerning the use of mouse models point to a lack of predictive validity and failure to translate into successful treatments (Onos et al., 2016). However, there are a variety of potential explanations for this disconnect between AD in mice and humans, including the fact that most mouse models of AD are maintained on only a single or a few genetic backgrounds. This includes backgrounds of mixed origins (Onos et al., 2016), which may confound interpretation of results. Previous studies have attempted to identify genes involved in the modification of AD symptoms in mice from mapping studies in F2 populations (Ryman et al., 2008; Sebastiani et al., 2006). However, these studies and individual strain-by-strain comparisons (Jackson et al., 2015; Sipe et al., 1993) have generally suffered from poor mapping resolution and/or an inability to pinpoint causative genes.

In order to identify specific genes involved in modifying resilience to AD symptoms in a more robust way, we developed the first AD transgenic mouse genetic reference panel. This panel, which we term the AD-BXDs, combines two well established resources: 1) the 5XFAD transgenic line that recapitulates various aspects of the human disease, including amyloid-β42 accumulation, cognitive deficits, and neuron loss (Oakley et al., 2006), and 2) the BXD genetic reference panel, the largest and best-characterized series of recombinant inbred strains derived from the two common inbred strains C57BL6/J (B6) and DBA2/J (D2) (Peirce et al., 2004). The BXD panel segregates more than 4.8 million sequence variants, including many in genes known to confer risk for AD (**Table S1**). This AD-BXD panel of F1 mice provides an ideal opportunity to monitor the phenotypic outcome of individuals harboring identical high-risk FAD mutations in human *APP* and *PSEN1* genes, raised in identical environments, but whose allelic contributions differ across the remainder of the genome. This approach parallels and complements searches for modifier genes that have been conducted in human patients, particularly modifiers of monogenic diseases caused by highly penetrant alleles similar to the 5XFAD transgene [i.e. the *CFTR* gene known to cause cystic fibrosis (Corvol et al., 2015)]. Overall, results presented here demonstrate the utility of the AD-BXD reference panel as a resource for the study of AD genetics and will likely provide critical insight into disease mechanisms.

## Results

### Genetic background modifies expressivity of FAD mutations

In order to evaluate the impact that genetic background has on the expressivity of human causal FAD mutations on behavioral and molecular phenotypes, we generated a panel of 26 genetically diverse F1 mouse strains with and without FAD mutations. Female B6 mice heterozygous for the dominant 5XFAD transgene (Oakley et al., 2006) were crossed to males from the BXD genetic reference panel (Peirce et al., 2004) to generate F1 progeny carrying the 5XFAD transgene (AD-BXDs, or carriers) or non-transgenic littermates (Ntg-BXDs, or non-carriers; **Figure 1A**). Working memory and body weight was monitored bi-monthly, and more in-depth phenotyping that included tests of motor function and anxiety was performed at both 6 and 14 months of age (**Figure 1B**). A subset of mice were subsequently tested for long-term spatial learning and memory function using a contextual fear conditioning (CFC) paradigm. This subset was immediately harvested following CFC testing, and tissue was collected for biobanking and later use, including RNA-sequencing and ELISAs as described below.

**Figure 1:**
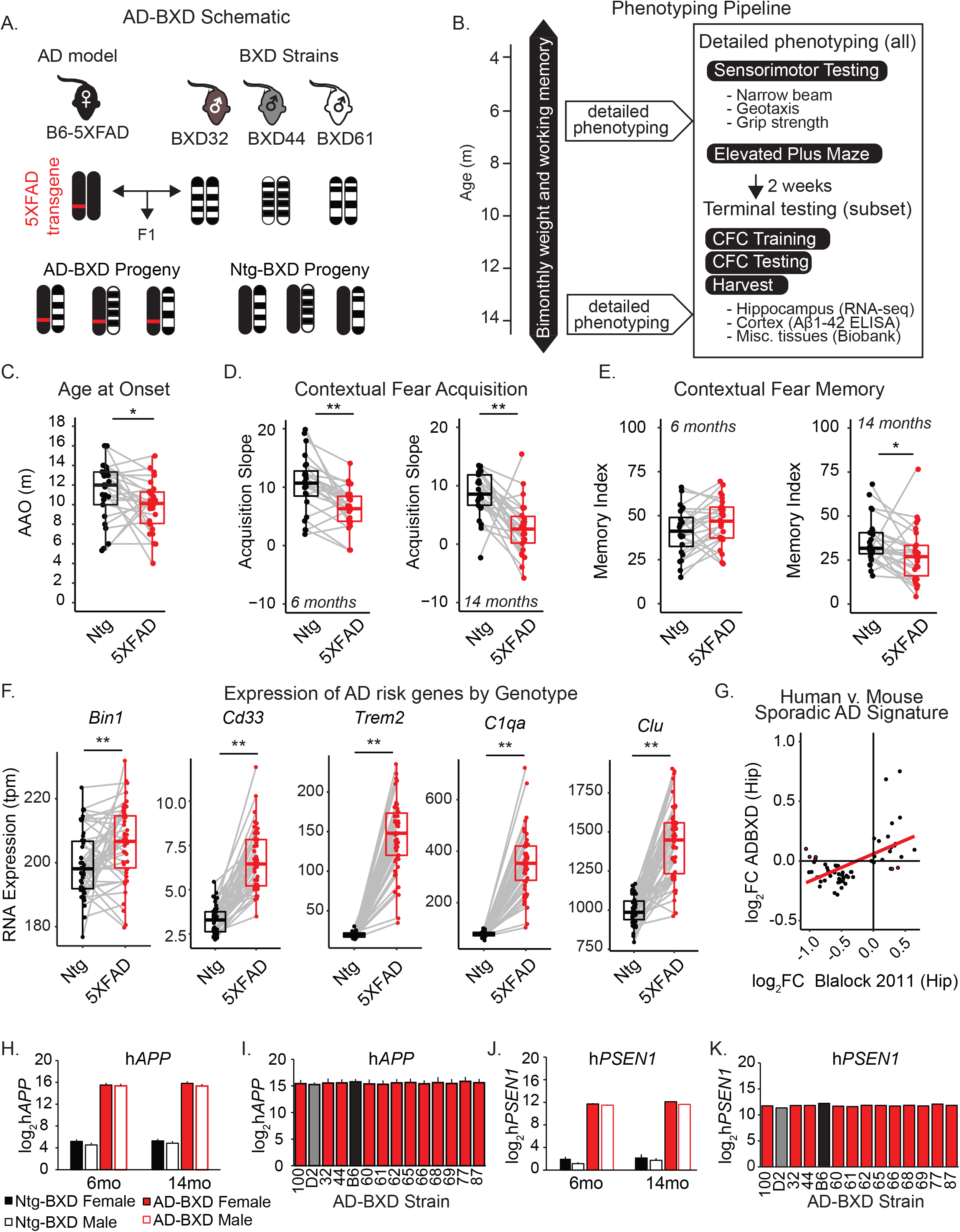
Genetic background modifies AD symptoms in a novel transgenic reference panel. A) Female C57BL/6J mice heterozygous for the 5XFAD transgene were bred to 26 BXD males to generate genetically diverse but isogenic F1 offspring. B) Body weight and working memory on the Y-maze was measured bi-monthly, and at 6 and 14 months more detailed phenotyping was carried out on a subset of mice. C) Onset of working memory deficits was significantly earlier in AD-BXDs compared to Ntg-BXD controls [as measured by average within-strain age at onset (AAO), AD-BXDs: n = 210 (109 females/101 males) across 24 strains vs Ntg-BXDs, n = 169 mice (106 female/63 male) across 25 strains, t(1,47) = 1.9, p = 0.03, one-tailed t-test]. D) AD-BXD mice exhibited poorer CFC acquisition at both 6 [left, t(1, 48) = 3.5, p = 0.001] and 14 [right, t(1, 49) = 5.0, p < 0.001] months. E) AD-BXD mice exhibit recall comparable to Ntg-BXDs during the CFM task at 6 months [left, t(1,48) = 1.5, p = 0.1] but are impaired by 14 months [right, t(1, 49) = 1.8, p = 0.04, one-tailed t-test]. For CFC, AD-BXD n = 26 strains, Ntg-BXD, n = 24-25 strains. F) Genes known to be associated with AD are differentially expressed in our panel. G) 50 out of 60 AD signature genes show concordant differential expression across mouse and human transcriptomes. The log2 fold change (FC) of differential gene expression between AD- and Ntg-BXDs is plotted on the y-axis, while the log2FC of gene expression between human AD patients and controls from Hargis and Blalock (2017) is plotted on the x-axis. Each point represents a single gene; discordant genes that change in opposite directions across mice and humans have been highlighted in red. H) RNA sequencing reads from the hippocampus of both AD and Ntg-BXDs were aligned to the human mutated sequence of APP and log transformed. Across the panel, there was a main effect of genotype [n = 133 (64 females/69 males across 14 strains), F(1, 132) = 1204, p < 0.001] but not of age, sex, or strain. I) Graph of hAPP expression across AD-BXD lines to illustrate there is no significant effect of background strain on transgene expression. J) Same analysis was done for RNA-sequencing reads aligned to the human mutated sequence of PSEN1. Across the panel, there was a main effect of genotype [n = 133 (64 females/69 males across 14 strains), F(1,132) = 1162, p < 0.001], but not of age, sex, or strain. K) Graph of hPSEN1 expression across the AD-BXD lines to illustrate there is no significant effect of background strain on transgene expression. For plots C-E, each point represents a strain average. For plot F, each point represents a strain/age/sex averaged sample. *p < 0.05 one-tailed t-test, **p < 0.05 two-tailed t-test. See also Figure S1 and Tables S1-3.

As expected, the 5XFAD transgene accelerated the age at onset (AAO) of working memory deficits in carriers relative to non-carriers (**Figure 1C**), and exacerbated CFC learning and memory deficits, particularly by 14 months (**Figure 1D-E**). In addition, the effect of the 5XFAD transgene significantly altered the expression of genes known to be misregulated in AD, particularly *Bin1, Cd33, Clu* (Karch et al., 2012), *Trem2* (Piccio et al., 2016), and *C1qa* (Hong et al., 2016), **Figure 1F**. However, the impact of causal FAD mutations on cognitive performance and risk gene expression varied widely depending on the specific background strain evaluated. In particular, the observed variation in AAO mirrors the variation in human patients reported by Ryman and colleagues (2014), suggesting our panel captures a portion of the phenotypic heterogeneity observed in human FAD patients. Heritability estimates comparing between-strain variance due to genetic diversity to total sample variance suggest there is a significant genetic component underlying observed variation, particularly on cognitive traits (**Table 1**). These results were not explained by an effect of strain, age, or sex on the transcription of the 5XFAD transgene itself as measured by alignment of RNA-sequencing reads to the mutated human *APP* or *PSEN1* sequence (**Figure 1H-K**). This suggests naturally occurring variants that segregate across the AD-BXDs play a significant role in determining susceptibility and/or resilience to changes in cognitive function and gene expression caused by high-risk FAD mutations.

**Table 1:** Heritability estimates for phenotypic traits in AD- and Ntg-BXDs. Heritability was determined by calculating the ratio of between-strain variance to total sample variance (variance due to both genetic and technical/environmental factors).

| Trait | Non-carriers (Ntg-BXD<sub>s</sub>) |  |  | Carriers (AD-BXD<sub>s</sub>) |  |  |
| --- | --- | --- | --- | --- | --- | --- |
| | Between strain variance | Total sample variance | Heritability ($h^2$) | Between strain variance | Total sample variance | Heritability ($h^2$) |
| Age at onset | 9.09 | 23.54 | 0.39* | 6.87 | 22.70 | 0.30* |
| 6m CFC Acquisition | 24.88 | 75.76 | 0.33* | 11.52 | 38.76 | 0.30* |
| 6m CFM | 196.63 | 537.13 | 0.37* | 164.58 | 527.95 | 0.31* |
| 14m CFC Acquisition | 14.69 | 65.02 | 0.23 | 18.52 | 65.21 | 0.28 |
| 14m CFM | 164.16 | 490.97 | 0.33* | 141.81 | 299.93 | 0.32* |
| 6m Grip strength | 0.09 | 0.28 | 0.32* | 0.13 | 0.32 | 0.40* |
| 14m Grip strength | 0.07 | 0.25 | 0.29 | 0.08 | 0.29 | 0.27 |
| 6m Narrow beam | 659.87 | 3011.14 | 0.22 | 561.91 | 3257.60 | 0.17 |
| 14m Narrow beam | 410.98 | 1827.03 | 0.22 | 1390.87 | 5696.10 | 0.24 |
| 6m Incline screen | 40.54 | 187.84 | 0.22 | 90.45 | 451.60 | 0.20 |
| 14m Incline screen | 79.75 | 445.05 | 0.18 | 307.64 | 1672.70 | 0.18 |
| 6m Sensorimotor composite | 0.33 | 2.51 | 0.13 | 1.12 | 4.99 | 0.22 |
| 14m Sensorimotor | 1.09 | 4.72 | 0.23 | 2.73 | 12.04 | 0.23 |
| 6m EPM % Time in Open Arms | 11.31 | 74.17 | 0.15 | 40.57 | 362.61 | 0.11 |
| 14m EPM % Time | 43.11 | 215.56 | 0.20 | 266.48 | 892.91 | 0.30* |

Abbreviations: m: months, CFC: contextual fear conditioning, CFM: contextual fear memory, EPM: Elevated Plus Maze \* heritability > 0.30 considered heritable

### AD-BXD transcriptome shows high concordance with sporadic late-onset AD signature

To evaluate the translational validity of incorporating naturally occurring genetic variation in AD models, we compared the expression of a set of 60 core genes previously defined as a human AD consensus signature (Hargis and Blalock, 2017) across a subset of AD-BXD and Ntg-BXD strains using RNA-sequencing of hippocampal tissue (**Table S2**). We observed high concordance between our mouse panel and human AD signatures – 83% (50 of 60) of genes in the human AD signature had fold changes in the same direction in our AD-BXD panel relative to Ntg-BXDs (**Figure 1G**). This effect replicated in 3 independent human datasets tested (**Figure S1**). Beyond this core set of 60 genes, our AD-BXD panel also displayed similarities to human AD in terms of functional enrichment of all significantly upregulated and downregulated genes (see Hargis and Blalock 2017 and **Table S3**). Pathways common to genes downregulated in human AD and the AD-BXDs include pathways related to the synapse (GABAergic, glutamatergic, and cholinergic synapse in mouse vs synapse part and synaptic vesicle in human). Pathways common to genes upregulated in both populations include cell adhesion (cell adhesion molecules in the mouse vs cell adhesion in human) and cell death (apoptosis in mouse vs cell death in human) (Hargis and Blalock, 2017). Overall, the incorporation of genetic diversity into a mouse model of AD resulted in a transcriptome profile that more closely matched human AD than previous AD models with limited genetic background variation as described by Hargis and Blalock (2017).

### Identification of genetic correlates of age at first symptom onset in the AD-BXD panel

To identify candidate mechanisms that influence susceptibility or resilience to FAD mutations, we sought to identify naturally occurring variants in the AD-BXDs specifically responsible for the observed heritable variation in AAO (**Fig 1C**). In order to define AAO, we used the Y-maze test of spontaneous alternation to longitudinally assess working memory in AD-BXD and Ntg-BXD mice. An animal was classified as impaired when performance dropped below chance levels (50%, indicating no memory for which arm of the maze was most recently visited) and the age at which impairment was first observed was termed the animal’s AAO. The average AAO for each AD-BXD strain varied widely [**Figure 2A**, n = 210 mice (109 female/101 male) across 26 strains - effect of strain, F(25,209) = 3.0, p = 0.04; no main effect of sex, p > 0.1], with susceptible strains exhibiting an AAO as early as 4 months and resilient strains maintaining cognitive function to the latest time point tested (**Figure 2B**). Notably, variation in AAO was not explained by differences in locomotor function or exploratory behavior (**Figure S2**).

**Figure 2:**
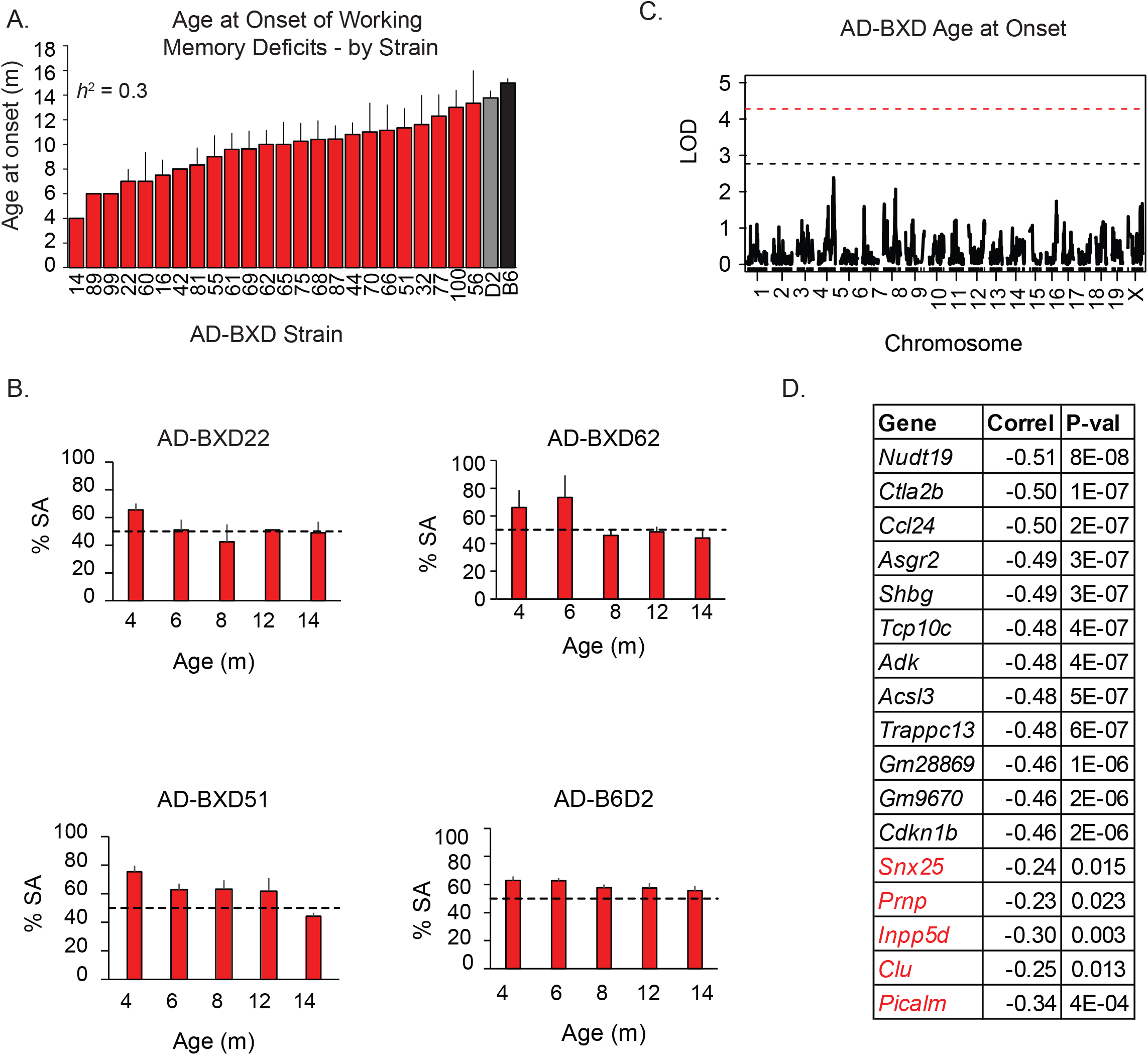
Age at which AD-BXD mice first exhibit working memory impairment exhibits heritable variation. A) Age at onset (AAO) of working memory deficits is significantly influenced by background strain [F(25,209) = 1.6, p = 0.04, n = 210 mice (109 female/101 male) across 26 strains]. Shown on the x-axis is the BXD strain used to create the AD-BXD line tested. B) Representative examples of working memory as measured by percent spontaneous alternation (%SA) scores across the lifespan. Dashed line represents chance performance (50%). C) Quantitative trait loci (QTL) mapping in r/qtl did not identify genomic loci regulating AAO. D) Several genes exhibited significant relationship with AAO including five genes previously associated with the AAO in human FAD populations (highlighted in red). See also Figure S2.

We first performed genetic mapping using r/qtl (Broman et al., 2003) in an attempt to identify specific loci (i.e. quantitative trait loci, QTLs) that are that are critical for modifying the AAO across the AD-BXDs. However, no significant QTL was identified for AAO (**Figure 2C**), and genotype at the peak marker on chromosome 4 explained only 12% of the variance. This result suggests multiple loci with relatively small effect sizes may regulate AAO. Consistent with this, we identified a relationship between AAO and the hippocampal expression of known modifiers of human FAD [**Figure 2D**, *Snx25* (Lee et al., 2015), *Prnp* (Dermaut et al., 2003), *Inpp5d, Clu, and Picalm* (Lambert et al., 2013) in red]. In addition, we identified twelve novel genes as associated with AAO after stringent Bonferroni correction for multiple comparisons (n = 21,215 genes, **Figure 2D in black**). As deficits in working memory performance in AD mouse models most closely parallels the earliest AD-related cognitive deficits detected in humans during the late preclinical phase of the disease (Webster et al., 2014), these gene candidates represent new targets to delay the onset of disease.

### Apoe *modifies contextual fear learning deficits in AD-BXDs*

Next, we sought to identify specific factors responsible for the observed variation in CFC acquisition in the AD-BXDs (**Fig 1D**). During CFC acquisition, a mouse that begins to associate the training context with the impending shock will exhibit increasing levels of freezing, a species-specific defensive response thought to indicate fear (Fanselow, 1984). The slope of the change in average freezing across the 4 post-shock intervals was used to derive an acquisition curve for each strain. Although the 5XFAD population was impaired relative to non-carrier littermates (**Figure 1D**) at both 6 and 14 months, the degree of this impairment varied widely [n = 355 mice (212 female/143 male) across 28 strains and 2 ages - effect of strain, F(27,354) = 2.1, p = 0.001, **Figure 3A-B**; no main effect of sex (p = 0.8)]. This variation was not associated with strain-specific variation in sensorimotor abilities, anxiety, or pain sensitivity as measured by post-shock reactivity (**Figure S3**). These results indicate the observed variation in contextual fear acquisition is regulated, in part, by genetic variants segregating across the AD-BXD panel.

**Figure 3:**
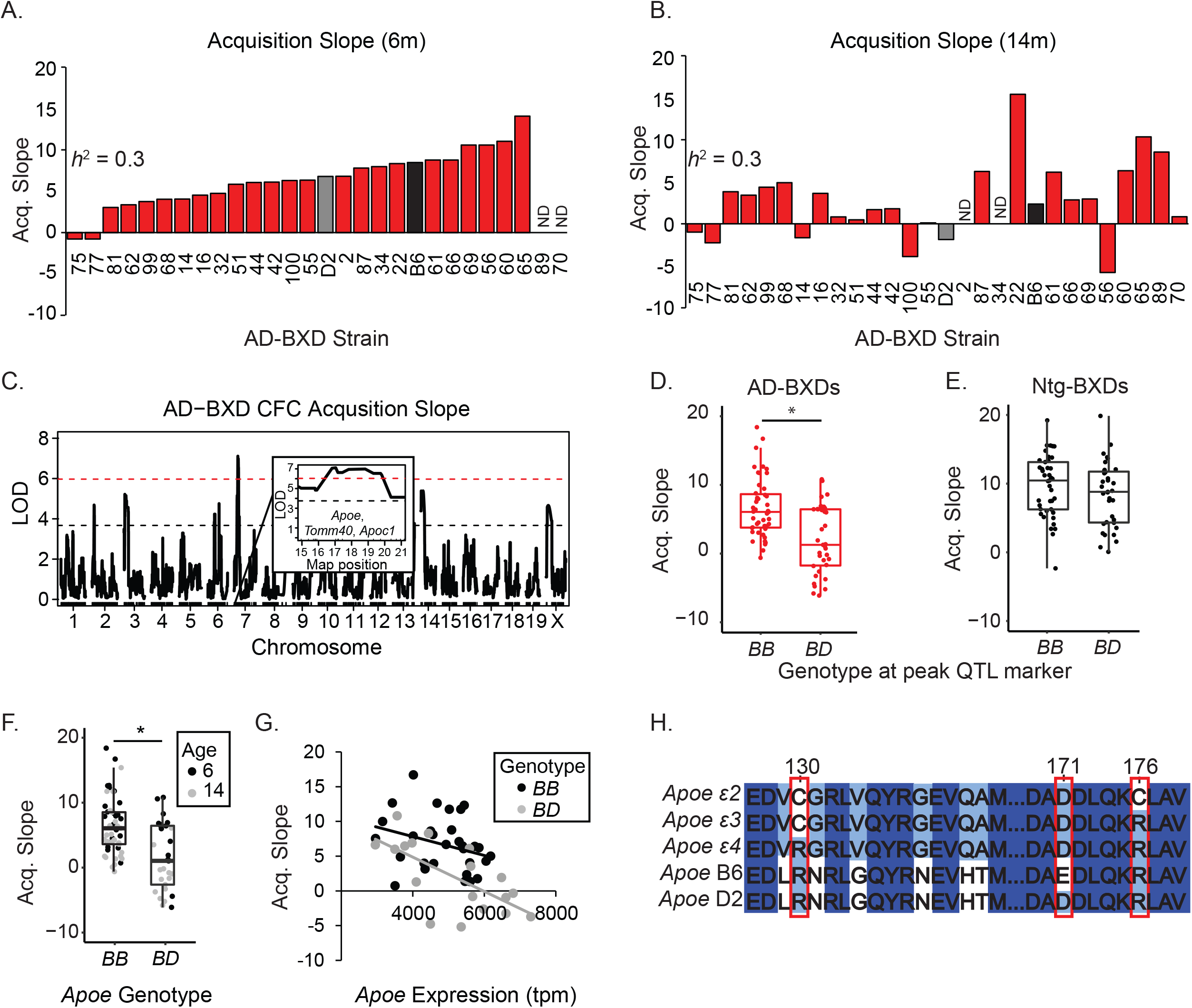
Conserved genetic mechanisms influence cognitive function in the AD-BXDs. A-B) For each strain, the average slope of the change in post-shock freezing following shocks 1-4 on training day was calculated. A significant effect of age [F(1,354) = 11.9, p = 0.001, n = 355 mice (212 female/143 male) across 28 strains and 2 ages] and background strain was detected [F(27,354) = 2.1, p = 0.001]. This variation was heritable at both 6 and 14 months (h2 = 0.3). C) Genetic interval mapping identified a significant QTL on chromosome 7 (1.5 LOD confidence interval = 15.90-20.38 Mb, LOD = 7.1). D) AD-BXD strains with a single D allele at the peak QTL marker exhibited significantly lower levels of CFC acquisition [t(1, 80) = 4.6, p < 0.001]. E) Genotype at peak QTL marker had no effect on acquisition slope in Ntg-BXDs [t(1,75) = 1.4, p = 0.15]. F) AD-BXD lines harboring the D Apoe allele performed significantly worse during contextual fear acquisition [t(1,80) = 4.5, p < 0.001]. G) Apoe expression is only significantly correlated with CFC acquisition when mice harbor a D allele at Apoe (genotype BD, r = -0.7, p < 0.01; genotype BB, r = -0.3, p = 0.10). H) The D allele harbors only a single SNP that causes the D2 sequence to match human ε4 variant at 3 critical amino acids (positions 130, 171, 174 in the full-length 317 amino acid human APOE). ND: no data. See also Figure S3 and Table S4.

Genetic mapping was performed using strain averages at both 6 and 14m, with age and sex included as covariates in a single QTL model. A significant QTL explaining 21% of the phenotypic variance was detected on chromosome 7 (1.5 LOD confidence interval = 15.90-20.38 Mb, LOD = 7.1, **Figure 3C**). AD-BXD strains harboring a *D* allele at this peak marker showed lower levels of acquisition [t(1, 80) = 4.6, p < 0.001, **Figure 3D**]. However, this effect was not observed when the peak marker was tested for association with acquisition in the Ntg-BXD population [t(1,75) = 1.4, p = 0.15, **Figure 3E**], demonstrating the effect of the identified QTL is specific to the AD-BXD population. This interval is syntenic to human 19q13.3 and contains approximately 100 genes with human orthologs and that contain either single nucleotide polymorphisms (SNPs), insertions/deletions (indels), or structural variants across our panel as annotated by the Sanger Mouse Genomes Project (Keane et al., 2011) – representing viable positional candidates that may contribute to observed variation in CFC acquisition. This included multiple genes previously shown to modify human individuals’ risk of AD, most notably *Apoe, Tomm40*, and *Apoc1*. All positional candidates with detectable levels of expression in the hippocampus (n = 70) were first filtered by correlation of hippocampal gene expression with CFC acquisition, as the hippocampus is critical for CFC (Holland and Bouton, 1999). Remaining after this analysis were 15 genes that passed stringent Bonferroni correction for association with CFC learning (**Table S4**). Of these genes, *Apoe* emerged as the candidate with strongest biological evidence for a causal involvement in phenotypic regulation. In addition to being the best-known risk factor for human AD, *Apoe* was the positional candidate most strongly related to genotype (log_2_FC Ntg-BXD vs AD-BXD = 0.8. adj. p = 7.1e-58) and age (log_2_FC 5XFAD adult vs 5XFAD aged = 0.4. adj. p = 6.8e-10) across our F1 population (**Table S4**). AD-BXD lines harboring a copy of the *D Apoe* allele performed significantly worse on acquisition of CFC regardless of age [t(1,80) = 4.5, p < 0.001, **Figure 3F**]. In addition, the correlation observed between *Apoe* expression and CFC acquisition (**Table S4**) appears to be driven by mice harboring the *D* allele, as expression was only significantly correlated with contextual fear acquisition when mice harbored the *D* allele (genotype *BD*, r = -0.7, p < 0.01; genotype BB, r = -0.3, p = 0.09; **Figure 3G**). This suggests that while more *Apoe* is overall detrimental to learning, this effect is exacerbated when the *D* allele of *Apoe* is present. Similar allele-specific results have been observed when over-expressing human *APOE* in transgenic mice (Bien-Ly et al., 2012). Interestingly, the *D* allele harbors only a single SNP when compared to the *B6* allele, and this missense mutation causes the D2 sequence to match the human ε4 variant more closely than the *B6* sequence (**Figure 3H**). As both alleles (ε4 and *D*) confer elevated risk for AD cognitive deficits, these results provide evidence that 1) the *D* allele of *Apoe* functionally recapitulates human *APOEε4* (Liu et al., 2013), and 2) shared genetic mechanisms confer susceptibility to AD in the AD-BXD and human populations. These results support the contention that discovery of novel genetic modifiers of AD symptoms using the AD-BXDs are likely to translate to humans.

### Susceptibility to AD memory deficits is regulated by genetic variants on chromosome 2

We next assessed the impact of the AD transgene in our panel on long-term contextual fear memory (CFM) as described previously (Neuner et al., 2015). AD-BXD mice exhibited impaired CFM by 14 months (**Figure 1E**), and the extent of variation in CFM at both age points varied significantly by strain [n = 355 mice (212 female/143 male) across 28 strains and 2 ages - effect of strain, F(27,354) = 3.6, p < 0.001, **Figure 4A-B**; no main effect of sex (p = 0.4)]. Variation in CFM was not association with strain-specific variation in gross sensorimotor abilities, anxiety, or pain sensitivity as measured by post-shock reactivity (**Figure S4**). Overall, results suggest variation in CFM is due primarily to strain-dependent differences in long-term memory and as a quantitative trait, is suitable for discovery of genomic loci associated with cognitive risk and resilience to AD mutations using QTL mapping.

**Figure 4:**
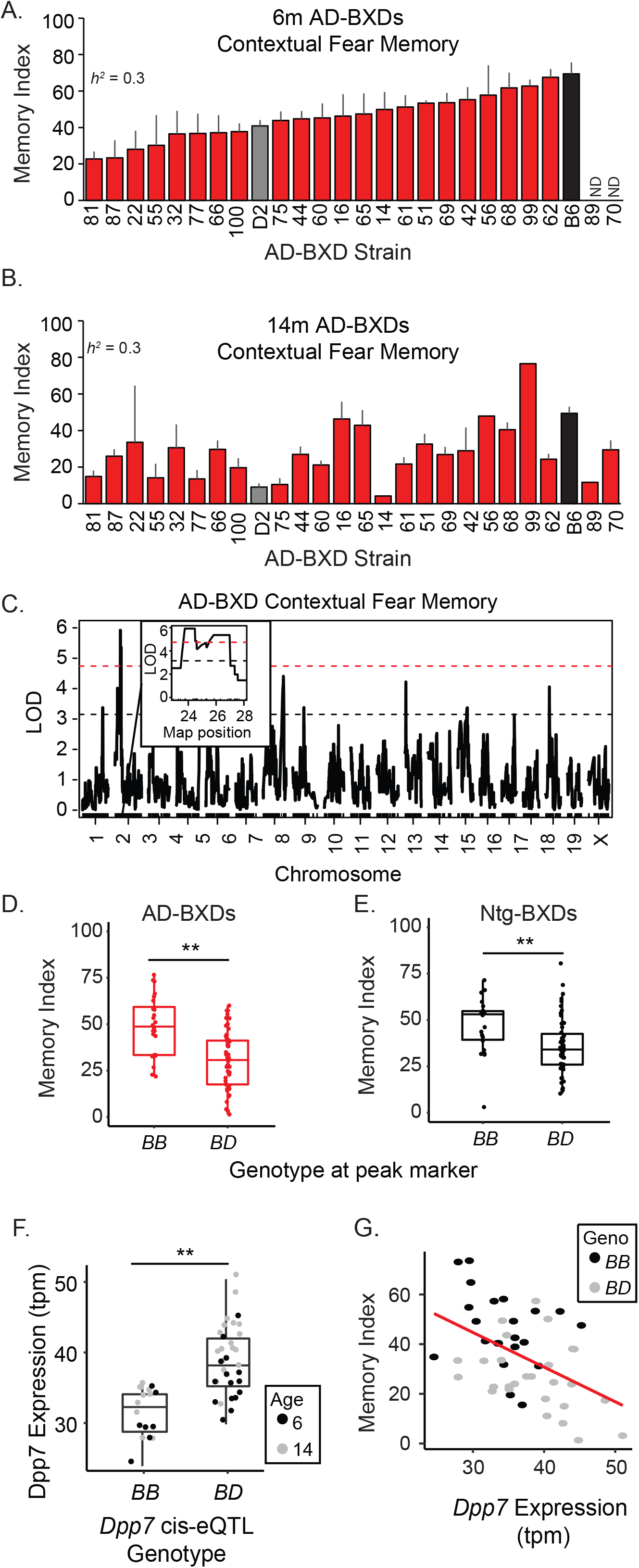
Contextual fear memory (CFM) is regulated by genetic variants on chromosome 2. A-B) Significant effects of age [F(1,354) = 34.9, p < 0.001] and strain [F(27,354) = 3.6, p < 0.001, n = 355 mice (212 female/143 male) across 28 strains] were detected in AD-BXD mice. This variation was heritable at both 6 and 14 months (h2 = 0.3). C) Genetic mapping identified a significant QTL on chromosome 2 (1.5 LOD confidence interval = 23.55-27.03 Mb, LOD = 5.9). D) AD-BXD lines harboring the D allele at the peak QTL marker exhibited significantly worse CFM [t(1,80) = 4.5, p < 0.001]. E) Genotype at peak marker also influenced CFM in Ntg-BXDs [t(1,75) = 3.0, p = 0.004]. E) Positional candidate Dpp7 is regulated by a local QTL on chromosome 2, and genotype at this cis-eQTL significantly alters Dpp7 expression [t(1, 46) = 5.4, p < 0.001]. F) Higher levels of Dpp7 expression are associated with poorer performance on CFM test (r = -0.47, p < 0.001). ND: no data. See also Figure S4 and Table S5.

QTL mapping was performed using strain averages at both 6 and 14 months, with age and sex included as covariates in a single QTL model. A significant QTL explaining 21% of the phenotypic variance was identified on chromosome 2 (1.5 LOD confidence interval = 23.55-27.03 Mb, LOD = 5.9 **Figure 4C**). In the AD-BXD panel, *D* alleles at the peak marker in this QTL were associated with lower levels of freezing and poorer cognitive performance [t(1,80) = 4.5, p < 0.001, **Figure 4D**). When tested for association with CFM in the Ntg-BXD panel, this marker also showed significant association with CFM [t(1,75) = 3.0, p = 0.004, **Figure 4E**]. However, this marker explained much less of the heritable variation in CFM in the Ntg-BXD panel (only 11% compared to 21% in the AD-BXD panel), suggesting that while this QTL may have broad relevance to CFM abilities, the effect is much more pronounced in the presence of the 5XFAD transgene. The QTL interval contained 62 positional candidates with naturally occurring genetic variants across our panel, six of which had gene expression significantly correlated with the CFM trait after stringent Bonferroni correction for multiple comparisons (**Table S5**). The expression of each these six candidates was mapped as separate quantitative traits in order to identify QTLs regulating observed variation in gene expression (expression QTL, or eQTL). A significant eQTL was identified only for *Dpp7* and overlapped the location of the gene itself [1.5 LOD confidence interval = 16.14 – 23.75, LOD = 9.0], indicating the expression of *Dpp7* is subject to local, or *cis*, regulation. As expected, genotype at the peak *cis* eQTL marker was significantly associated with gene expression, with one copy of the *D* allele associated with much higher levels of hippocampal *Dpp7* expression [t(1, 46) = 5.4, p < 0.001, **Figure 4F**]. As expected considering the association of *D* alleles with poorer cognitive performance, higher levels of hippocampal *Dpp7* expression were significantly associated with poorer CFM (r = -0.47, p < 0.001, **Figure 4G**). Taken together, these data suggest a role for *Dpp7* as a negative regulator of CFM in the AD-BXD panel and a candidate modifier of susceptibility to AD.

### AD-BXD panel exhibits heritable variation in amyloid pathology attributable to genetic variants on chromosome 3

In addition to deficits in cognitive function, one of the hallmarks of AD is accumulation of toxic Aβ 1-42 (Aβ42), thought to be an initiating factor in a cascade of symptoms eventually leading to neuron loss and dementia (Hardy and Higgins, 1992). To measure Aβ42 levels across the panel, brain extracts from 24 distinct AD-BXD lines were assayed in duplicate on Aβ42-specific sandwich ELISA (n = 154 mice (89 female, 65 male) across 23 strains and 2 ages, **Figure 5A-B**). Variation in Aβ42 levels were heritable (*h*^2^ > 0.30), and increased with age [F(1,153) = 132.8, p < 0.001]. A significant main effect of strain was observed [F(22,153) = 2.1, p = 0.01], indicating that genetic background significantly modified the Aβ42 levels across the panel. No main effect of sex was observed [F(1, 153) = 1.4, p = 0.25], in contrast to a previous report using a single genetic background suggesting the 5XFAD model preferentially affect females (Sadleir et al., 2015).

**Figure 5:**
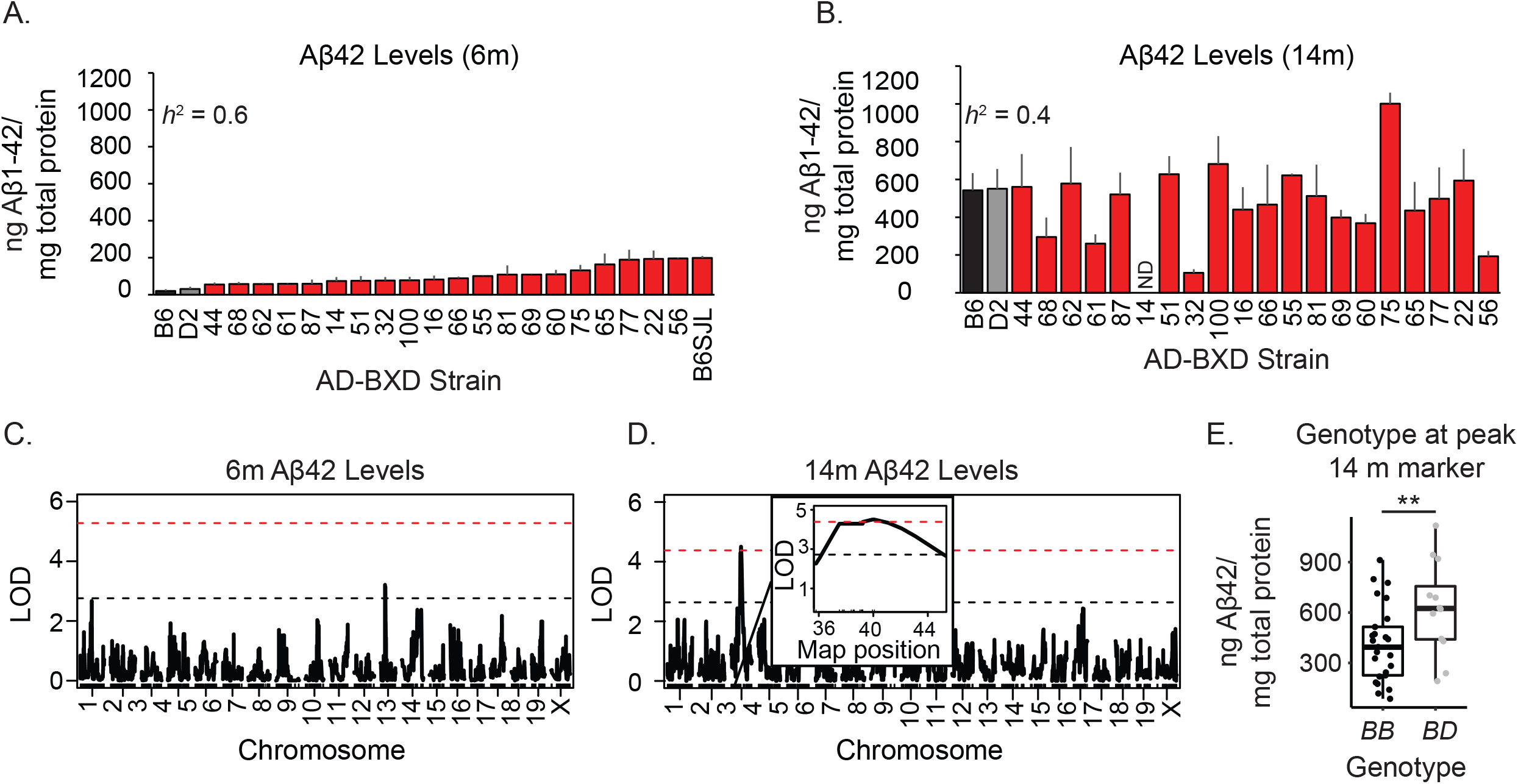
QTL on chromosome 3 regulates variation in amyloid pathology across the AD-BXDs. A-B) Aβ42 drastically increased from 6 to 14m [effect of age, F(1,153) = 132.8, p < 0.001, n = 154 mice (89 female, 65 male) across 23 genotypes] but varied significantly by background strain [effect of strain, F(22,153) = 2.1, p = 0.01]. C) No significant QTL was identified for Aβ42 levels at 6 months. D) A significant QTL on chromosome 3 was detected for Aβ42 load at 14 months (1.5 LOD confidence interval = Chr 3: 36.08-45.08 Mb, LOD = 4.5). E) D alleles at the peak marker contribute higher Aβ42 levels across the panel [t(1,35) = 2.5, p = 0.02]. ND: no data. See also Figure S5 and Table S6.

In order to identify genetic factors involved in regulating Aβ42, r/qtl was used to perform genetic mapping. While no significant QTLs were detected for Aβ42 load at 6 months (**Figure 5C**), a significant QTL on chromosome 3 explaining 15% of the phenotypic variance was detected for Aβ42 load at 14 months (1.5 LOD confidence interval = Chr 3: 36.08-45.08 Mb, LOD = 4.5, **Figure 5D**). *D* alleles at the peak marker associated with higher Aβ42 levels across the panel (**Figure 5E**). Within this QTL, there were 22 positional candidates with sequence differences between B6 and D2, each of which represent viable positional candidates that may directly contribute to the variation in Aβ42 levels (**Table S6**). No candidate exhibited significant correlation between hippocampal gene expression and amyloid levels, suggesting to us that the identified variant may exert its effect through alteration of protein function rather than transcript expression. Of the 20 positional candidates, six candidates contained both SNPs and indels predicted to have a high functional impact on either transcript or protein structure according to Ensembl Variant Effect Predictor (McLaren et al., 2016). Three of these six genes, fibroblast growth factor 2 (*Fgf2*) (Donnini et al., 2006; Katsouri et al., 2015), cyclin A2 (*Ccna2*) (Chandrasekaran and Bonchev, 2016), and interleukin 2 (*Il2*) (Alves et al., 2017) have previously been associated with Aβ levels, suggesting variants in one or multiple genes in this region may influence amyloid load. Of the remaining 3 genes that had not been previously implicated as involved in amyloid accumulation, *Trpc3* emerged as the gene with the most biological evidence for a potential functional role in AD.

### *Targeted knockdown of positional candidate* Trpc3 *reduces Aβ42 load and AD-related cognitive symptoms*

*Trpc3* is a member of the transient receptor potential channel family and is permeable to cations including calcium (Dietrich et al., 2005). Misregulation of calcium signaling has previously been implicated in the pathogenesis of AD (LaFerla, 2002), and *Trpc3* itself has recently been implicated in neuronal excitability and cognitive function in adult mice (Neuner et al., 2015). In addition, *Trpc3* function has been shown to be sensitive to cellular cholesterol (Graziani et al., 2006), a pathway closely linked to AD by GWAS hits such as *APOE, CLU*, and *ABCA7* (Karch and Goate, 2015). Across the AD-BXDs, *Trpc3* contains both a sequence variant and an insertion in a predicted splice region, further strengthening a possible role for *Trpc3* in in cognitive deficits and amyloid processing. Notably, the insertion occurs in exon 10 of *Trpc3* (NM_019510), and a calmodulin/IP_3_R binding site within exons 9 and 10 has previously been shown to modulate TRPC3 activation (Zhang et al., 2001). Given that antibodies directed against beta-amyloid have not resulted in disease-modifying treatments, and beta-amyloid does not correlate strongly with cognition in either humans or our AD-BXD panel (**Figure S5**), we hypothesized that targeting a putative modulator of amyloid pathology that also has cognitive function-enhancing capabilities (i.e. *Trpc3*) may provide an added benefit to susceptible strains by reducing pathology and increasing neuronal excitability.

Since TRPC3 protein is increased in hippocampus from 5XFAD mice compared to Ntg controls [**Figure 6A**, n = 4/grp, t(1,6) = 3.7, p = 0.01], we injected our previously validated AAV9 viral vector containing either shRNA targeting *Trpc3* (shRNA-Trpc3) or a scrambled shRNA control (shRNA-Ctrl) (Neuner et al., 2015) directly in the dorsal hippocampus of presymptomatic male 4 month-old 5XFAD-B6SJL mice (Oakley et al., 2006). This strain was chosen as it a susceptible strain with robust amyloid deposition when compared to mice from our AD-BXD panel (**Figure 5A**). The mice were aged to 9 months, a time point at which a majority of the population exhibits both amyloid accumulation and memory deficits (Kaczorowski et al., 2011; Oakley et al., 2006), and then working memory and CFM was assessed (**Figure 6B**).

**Figure 6:**
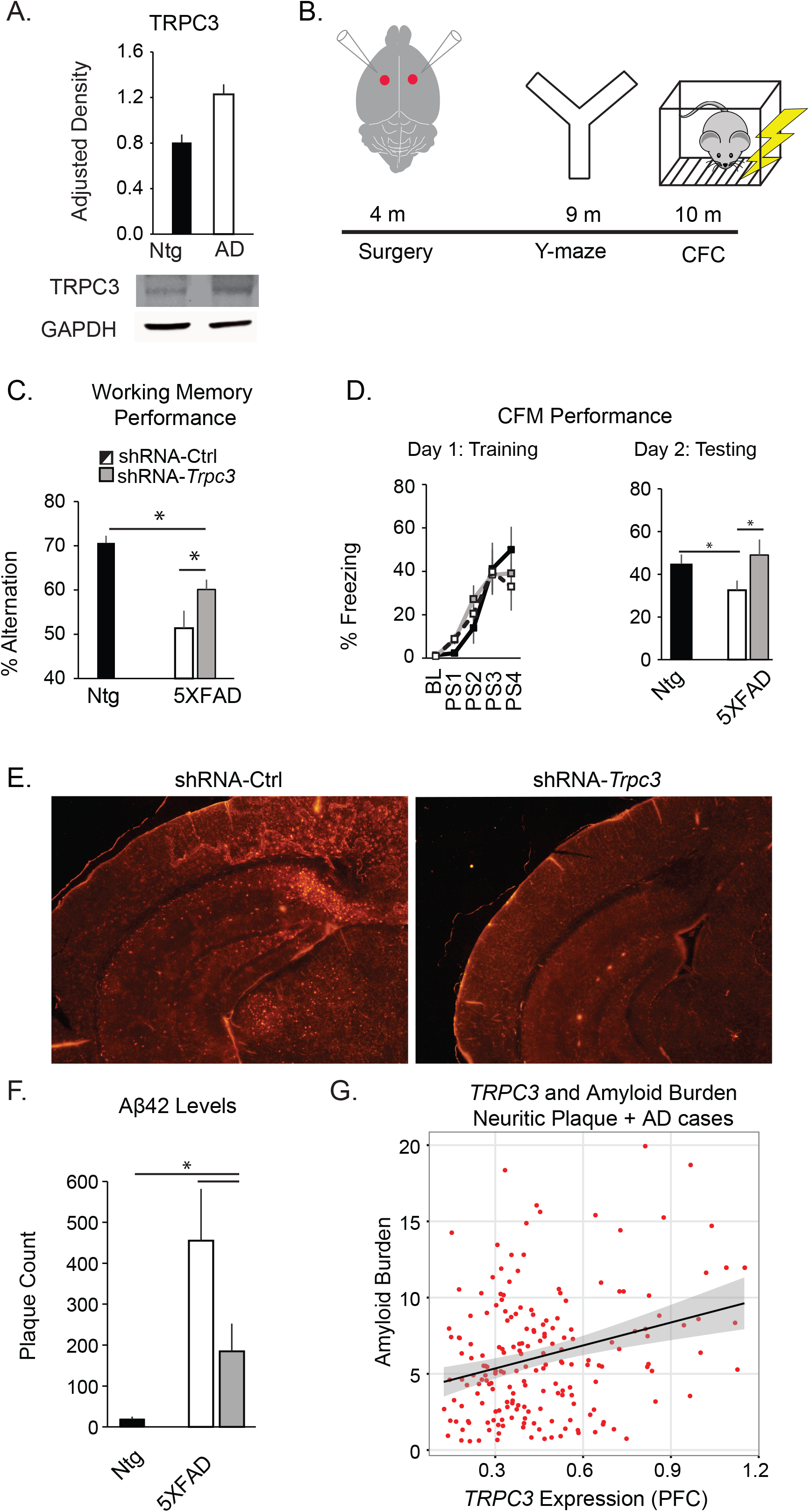
Knockdown of positional candidate Trpc3 delays AD symptoms in a mouse model of AD. A) TRPC3 protein is increased in the hippocampus of 5XFAD mice as measured by Western blot [n = 4/grp, t(1,6) = 3.74, p = 0.01]. B) AAV9 encoding either shRNA targeting Trpc3 (shRNA-Trpc3) or a scrambled control shRNA (shRNA-Ctrl) was delivered bilaterally into the hippocampus of presymptomatic 4 month-old 5XFAD mice. Working memory was assessed on the y-maze at 9m while CFM was assessed at 10m. C) There was a significant effect of group on working memory [F(2, 28) = 10.6, p < 0.001], with 5XFAD shRNA-Ctrl mice exhibiting impairment relative to both Ntg shRNA-Ctrl mice [post-hoc t-test, t(1,18) = 4.4, p < 0.001] and 5XFAD shRNA-Trpc3 mice [t(1,18) = 2.2, p = 0.04]. D) Left, all mice exhibited similar levels of freezing after the final training shock, indicating comparable acquisition across groups [F(2,26) = 1.1, p > 0.3]. Right, there was a significant effect of group on CFM [F(2,26) = 3.4, p = 0.05]. 5XFAD shRNA-Ctrl mice exhibited impairment relative to both Ntg shRNA-Ctrl mice [t(1,17) = 2.3, p = 0.03] and 5XFAD shRNA-Trpc3 mice [t(1,18) = 2.1, p = 0.047], who performed comparably (p > 0.05). E) Left, representative 2x images of coronal brain sections from 10m 5XFAD mice. The levels function in Photoshop was used identically across all images to increase visibility. There was a significant effect of group on number of Aβ42 immunoreactive plaques in the hippocampus and cortex [F(2,11) = 5.5, p = 0.03]. 5XFAD shRNA-Trpc3 mice showed a decrease in the number of plaques in the hippocampus and cortex compared to 5XFAD shRNA-Ctrl mice [one-tailed t-test, t(1,7) = 2.0, p = 0.04] and were not significantly different than Ntg mice [t(1,6) = 1.8, p = 0.12). G) A significant association was observed between TRPC3 expression in the prefrontal cortex and a measure of brain-wide amyloid burden in neuritic plaque positive human AD cases (p = 0.0005). See also Figures S6 and S7.

As expected, 5XFAD mice injected with shRNA-Ctrl performed significantly worse than Ntg littermates injected with shRNA-Ctrl on both working memory [**Figure 6C** - effect of group, F(2, 28) = 10.6, p < 0.001; post-hoc 5XFAD shRNA-Ctrl vs Ntg shRNA-Ctrl: t(1,18) = 4.4, p < 0.001] and CFM tests [**Figure 6D** - effect of group, F(2,26) = 3.4, p = 0.05; post-hoc 5XFAD shRNA-Ctrl vs Ntg shRNA-Ctrl: t(1,17) = 2.3, p = 0.03]. Consistent with our hypothesis, 5XFAD mice that had received shRNA-Trpc3 performed better than 5XFAD mice receiving control injections [post-hoc 5XFAD shRNA-Trpc3 vs 5XFAD shRNA-Ctrl, p < 0.05 on both tasks] and were statistically indistinguishable from controls on the CFM task (post-hoc 5XFAD shRNA-Trpc3 vs Ntg shRNA-Ctrl p > 0.05). Importantly, no significant effects of group on total distance traveled or arms entered in the y-maze, or baseline freezing during CFC training, were observed indicating no difference between groups on measures of total activity or anxiety (**Figure S6**). Similar effects were observed in non-transgenic mice, demonstrating a general effect of *Trpc3* knockdown on cognitive aging (**Figure S7**).

Finally, we investigated the effects of *Trpc3* knockdown on accumulation of Aβ42. Mice were harvested at 10 months and one hemisphere of each brain region was fixed in 4% PFA. Immunohistochemistry for Aβ42 was performed and number of plaques counted using Image J’s particle analysis software (Hurtado et al., 2010). 5XFAD shRNA-Ctrl mice exhibited a robust increase in the total number of plaques observed in the cortex and hippocampus (**Figure 6E**). In contrast, 5XFAD mice treated with shRNA-*Trpc3* showed a decrease in the number of plaques [effect of group: F(2,11) = 5.5, p = 0.03; post-hoc one-tailed t-test 5XFAD shRNA-Ctrl vs 5XFAD shRNA-*Trpc3*: t(1,7) = 2.0, p = 0.04] and were not significantly different than Ntg mice [post-hoc t-test 5XAFD shRNA-*Trpc3* vs Ntg shRNA-Ctrl: t(1,6) = 1.8, p = 0.12]. These results demonstrate that in addition to a general role in cognitive aging, *Trpc3* plays a disease-specific role in the regulation of amyloid levels in AD.

To assess the translational relevance of this finding, we next evaluated the relationship between TRPC3 and human AD. Using ROS/MAP data, a significant association was identified between *TRPC3* expression in the prefrontal cortex of neuritic plaque positive AD cases and a brain-wide measure of amyloid burden (see Methods), even after adjusting for age at death and sex (p = 0.0005, **Figure 6G**). As in our mouse panel, higher levels of *TRPC3* were associated with an increased amyloid burden. In contrast, no association was observed between *TRPC3* expression and neurofibrillary tangles, cerebral amyloid angiopathy, or Lewy bodies (p > 0.05). In addition, publically available data was mined for evidence of *TRPC3* association with human AD. Of three SNPs annotated to *TRPC3* in the International Genomics of Alzheimer’s Project (IGAP) dataset (Lambert et al., 2013), one SNP displayed a nominal association with AD (rs114991240, uncorrected p = 0.03). Finally, *TRPC3* appears in the same module as known AD risk genes *APOE, CLU*, and *DSG2* in a gene regulatory network constructed from post-mortem brain tissue from LOAD patients and cognitively normal controls (Zhang et al., 2013), suggesting its role in a larger regulatory network may influence risk of AD. Together, these results suggest that while variants in *TRPC3* itself may not play a highly significant role in regulating risk of AD in human populations, mechanisms and pathways in which *TRPC3* is involved (e.g. neuronal excitability, cholesterol metabolism, amyloid production and clearance) are important for modulating risk of AD. Overall, results here implicate *Trpc3* for the first time in regulation of AD pathogenesis and demonstrate the ability to transition from candidate gene identified by QTL mapping to functional validation in an *in vivo* mouse model and translational evaluation using human datasets.

Discussion

*Transgenic reference panel allows for new approach to study AD genetics*

It has long been recognized in humans that Alzheimer’s disease is a complex and polygenic disease, likely influenced by multiple variants, some with relatively small effect sizes (Lambert et al., 2013). Here, we introduce the first genetically diverse population of AD mice - the AD-BXDs - as a useful model of human AD by demonstrating a high level of concordance between the AD-BXDs and human AD, both phenotypically and at the level of hippocampal transcriptome profiles. We further validated the AD-BXDs at the genetic level by identifying *Apoe* as a positional candidate located in a genomic interval significantly associated with cognitive deficits, and demonstrated the ‘risk’ allele in our population most closely matches the deleterious ε4 allele in humans. These results also validate CFC as a behavioral assay with predictive validity relative to human GWAS studies. Finally, using a combination of forward genetics and *in vivo* functional validation, we identify *Trpc3* as a modulator of both age-related cognitive deficits and neuropathological accumulation of Aβ42 in AD. While multiple genes likely regulate cognitive function and Aβ42 accumulation in AD, the identification and validation of *Trpc3* provides a novel therapeutic target and mechanistic pathway to be pursued in future studies.

Work presented here expands on several previous studies in mouse models that have attempted to identify precise genes involved in the modulation of AD pathology phenotypes (Ryman et al., 2008; Sebastiani et al., 2006). However, in both cases, genetic intervals identified were too large to pinpoint causative variants. We utilized the BXD GRP for several reasons, including the fact that recombination intervals are much smaller than those observed in F2 populations, allowing for the identification of narrower QTLs than observed previously (Peirce et al., 2004). Since each BXD line is fully inbred, each F1 AD-BXD line studied here can be reproduced across time and laboratories, maximizing the utility of our characterization of AD-BXD lines as either susceptible or resilient to AD. Our F1 strategy is also advantageous as it decreases the number of homozygous alleles in a panel and enhances hybrid vigor (Charlesworth and Willis, 2009). Lastly, due to this reduction of homozygous alleles, allele-specific expression of a given gene can be monitored and quantified in a systematic way (Pandey and Williams, 2015).

### The role of modifier genes in normal aging and AD

The creation of the AD-BXD panel enables genetic mapping to identify true modifier alleles that influence the onset and severity of disease. However, interpretation of these modifiers in the context of normal aging is complex, as the extent to which genetic mechanisms underlying normal cognitive aging and AD overlap is still unclear. In principle, modifier alleles of interest could act in one of two ways: 1) as general modifier alleles that contribute to a phenotype regardless of disease status, or 2) specific AD modifiers that exhibit some type of epistatic relationship with the 5XFAD transgene and show no effect on phenotype in Ntg-BXD mice. Interestingly, we see examples of both instances in our results, suggesting both mechanisms are at play in our panel. For example, the peak marker in the QTL for CFM on chromosome 2 showed a significant association with CFM abilities in both AD- and Ntg-BXD mice (**Figure 4D-E**). However, in Ntg-BXD mice, this locus explained a smaller proportion of the variance (11%) compared to AD-BXD mice (21%) and would not have been detected by standard QTL mapping. Thus, the inclusion of the 5XFAD transgene as a sensitizer enabled the identification of a QTL that likely plays a small role in CFM abilities regardless of genotype, but which would have failed to be identified in studies of normal aging. Genes located in this interval likely represent broadly applicable therapeutic targets due to their role in both normal aging and AD. As an example of the second potential modifier mechanism, the QTL for CFC acquisition played no discernable role in regulating learning abilities in the Ntg-BXD population, suggesting this locus played a genotype-specific role and may act as an epistatic modifier of the 5XFAD transgene. These two examples highlight the particularly complex relationship between normal aging and Alzheimer’s disease. It is also interesting to note that, as in humans, our identified QTLs do not explain all of the variation in their respective traits that are attributable to genetic factors. While this can be improved by increasing mapping power, our results and the lack of a QTL in AAO mapping further illustrates the point that multiple small effect variants with different roles (e.g additive, epistatic) in disease susceptibility contribute to a portion of the heritability of this complex disease.

### Importance of genetic background in choosing a mouse model

Decades of research using AD mouse models have failed to translate into successful treatments for a variety of reasons, many of which have been discussed previously (Onos et al., 2016). Our results demonstrate that genetic variation profoundly affects the expressivity of the 5XFAD transgene on both cognitive and pathological traits. Thus, we conclude that a significant contributor to prior negative outcomes may be the lack of genetic diversity. Moreover, we demonstrate that the C57BL/6J genetic background harbors resilience factors that attenuate the impact of 5XFAD mutations on CFM (**Figure 4B**), despite moderate-to-high levels of Aβ42 (**Figure 5B**). While this suggests the C57BL/6J mouse may be a valuable model in which to study resilience to AD-related neuropathologies, transgenic AD models created on the C57BL/6J background may pose problems for screening drug targets or understanding mechanisms that drive *risk* of AD. Along with other genetically diverse models, the AD-BXD panel may provide new opportunities for preclinical screening that is directly beneficial to human patients. Since the 5XFAD transgene is not traditionally thought to induce significant tau pathology (Oakley et al., 2006), we did not evaluate tau pathology in the AD-BXDs. However, naturally occurring variants in*Mapt* [see Sanger Mouse Genomes Project (Keane et al., 2011)] segregate across our AD-BXD panel and may influence the production of hyperphosphorylated tau (and ultimately, neurofibrillary tangles). As tau and amyloid rarely occur together in mouse models, these strains (if identified) would both provide ideal models for preclinical screens in future studies and contribute to our understanding of the genetic variants modifying production and/or clearance of hyperphosphorylated tau. In addition, as tau has been seen to be more strongly associated with cognitive function in AD than other pathologies (Brier et al., 2016), the AD-BXD panel may provide new opportunities to study the relationship between tau, amyloid, and cognitive function in the context of genetic diversity.

### Trpc3 *as a novel modulator of AD resilience*

In order to contribute new understanding of mechanisms underlying resilience to disease, we pursued functional validation of *Trpc3*, an ion channel that had not previously been associated with AD, but for which we had strong biological evidence supporting a potential role in disease. *Trpc3* is known to be permeable to calcium, and misregulation of calcium signaling is known to be one of the earliest events that occur in disease pathogenesis. Calcium regulation has also been shown to directly modulate Aβ production, thereby providing a mechanistic link between *Trpc3* channels and Aβ42 modulation (LaFerla, 2002). In addition, our lab has previously implicated *Trpc3* as causally involved in the regulation of neuronal excitability and cognitive function, providing a biological link between *Trpc3* channels and cognitive symptoms of AD (Neuner et al., 2015). It was our hypothesis that targeting a gene that is both a known cognitive enhancer and putative mediator of Aβ42 levels would likely provide a double benefit to AD carriers by better allowing neurons to participate in networks critical for learning. Finally, a close family member of *Trpc3, Trpc6*, was recently shown to directly interact with amyloid precursor protein (APP), leading to a change in processing by gamma-secretase and a reduction in overall Aβ levels (Wang et al., 2015). Additional work is needed to determine whether *Trpc3* mediates its effects on AD-related symptoms through regulation of calcium, neuronal excitability, or through a direct interaction with APP reminiscent of *Trpc6*. Regardless, the discovery that decreasing expression of *Trpc3* in either normal aging or an aggressive mouse model of AD is sufficient to delay the onset of both cognitive and disease-specific pathological symptoms greatly contributes to our understanding of AD genetics, how they relate to mechanisms underlying normal aging, and provides an additional target on which to focus future studies.

One drawback of our current study is the lack of exhaustive functional testing of each positional candidate located in the QTL identified for Aβ42. It is likely that additional genes play a role, either independently or via an interaction with *Trpc3*, in determining severity of AD neuropathology. This idea is supported by the fact that positional candidates *Fgf2* and *Il2* have independently been associated with amyloid levels in model systems (Alves et al., 2017; Katsouri et al., 2015). In addition, the QTL identified in our AD-BXD panel does not overlap with those broad regions associated with pathology previously (Ryman et al., 2008; Sebastiani et al., 2006). While there are a variety of reasons why this may be the case, including the use of different experimental populations and quantitative traits, this phenomenon is likely best explained by the fact that AD is an incredibly complex disease (Rollo et al., 2016), accompanied by complex endophenotypes such as Aβ42 accumulation, and is likely to be mediated by multiple genes with small effect sizes. Therefore, continued expansion of the AD-BXD panel and/or related resources with even greater genetic diversity and power may enable discovery of many additional genes and mechanisms.

### Conclusions and future directions

The ultimate goal of mouse studies relating to AD is the eventual translation of identified candidates into viable human therapeutics or biomarkers of disease. Our results suggest the AD-BXD panel is a valuable resource to do just that, as we demonstrate high levels of overlap between the AD-BXDs and human AD at the phenotypic, transcriptomic, and genetic level. In addition, we demonstrate genotype at *Trpc3* is important for in mouse models determining resilience to AD, particularly AD neuropathology. In addition to utility as a potential biomarker, *Trpc3* may be able to be targeted mechanistically to treat disease. The translational relevance of this idea is supported by the association identified here between levels of *TRPC3* in human patients and a measure of brain-wide amyloid burden. Currently, selective antagonists for *Trpc3* are commercially available (Kiyonaka et al., 2009), however they generally have poor bioavailability and contain potentially toxic moieties. Current work is aimed at adapting these antagonists to extend half-life and reduce the potential side effects of administration, with the goal of easily manipulating *Trpc3* function *in vivo*. Although there is a distinct possibility *Trpc3* may not provide a direct translational target to human populations, results generated here and in future studies examining the mechanistic role of *Trpc3* in AD may provide important insight into pathways important for Aβ42 generation. Future therapeutics may therefore focus on *Trpc3* or targets/cellular mechanisms that exist downstream of *Trpc3* itself.

## Author Contributions

SMN and CCK conceived of the experiments. SMN, KMSO, and CCK designed the experiments and analyses, and wrote the manuscript. SMN conducted the behavioral and QTL mapping experiments, while RH and MJH conducted the RNA sequencing experiments and assisted in analyzing the data and interpretation of results. SMN and TH conducted the human validation analyses. DAB, JAS, and PLDJ contributed human data and assisted with analyses and interpretation of results. All authors reviewed and approved of the final manuscript.

## Acknowledgements

This work was supported by the BrightFocus Foundation (A2016397S to CCK), the National Institute on Aging (R01AG054180, R01AG057914 to CCK, F31AG050357 to SMN, K01AG049164 to TJH), and the National Institute of Diabetes and Digestive and Kidney Diseases (R01DK102918 to KMSO). Study data for the human validation studies were provided by the Rush Alzheimer’s Disease Center, Rush University Medical Center, Chicago. Data collection was supported through NIA grants P30AG10161, RF1AG15819, and R01AG17917 to DAB, R01AG36836 to PLD, R01AG30146, U01AG32984, U01AG46152, the Illinois Department of Public Health, and the Translational Genomics Research Institute. The authors thank Dr. Lynda Wilmott and Thomas Shapaker for collection of behavioral data and Matthew de Both and Ashley Siniard for assistance with RNA-sequencing.

## Methods

### CONTACT FOR REAGENT AND RESOURCE SHARING

Further information and requests for resources and reagents should be directed to and will be fulfilled by the Lead Contact, Catherine C. Kaczorowski.

### EXPERIMENTAL MODEL AND SUBJECT DETAILS

Female congenic C57BL/6J mice hemizygous for the dominant 5XFAD transgene (Oakley et al., 2006), which consists of 5 human mutations known to cause familial AD [three in amyloid precursor protein (APP; Swedish: K670N, M671L, Florida: I716V, and London: V717I) and two in presenilin 1 (*PSEN1*; M146L and L286V)], were obtained from The Jackson Laboratory (JAX MMRRC Stock No: 34848-JAX). These mice were bred with 26 males from a set of genetically diverse recombinant inbred strains from the well-established BXD genetic reference panel (Peirce et al., 2004) (**Fig 1A**). Lists of the BXD strains used here and corresponding genotypes (mm9) are available as **Table S7**. By selecting the same maternal background strain (i.e. 5XFAD-C57BL/6J) across the panel for cross with male BXD strains, we were able to introduce variants in the nuclear DNA, hold the mitochondrial genome constant, and control for strain-specific differences in maternal behavior on offspring behavior. The F1 progeny resulting from this B6-5XFAD by BXD cross are isogenic recombinant inbred backcross mice, each harboring one maternally derived *B* allele and either a *B* or *D* paternally derived allele at any given genomic locus. As expected from a Mendelian pattern of inheritance, ~50% of these F1 mice carry the 5XFAD transgene (termed AD-BXDs, or carriers) and ~50% are non-transgenic (Ntg) littermate controls referred to Ntg-BXDs (or non-carriers). Male and female offspring were group housed (2-5 per cage) and maintained on a 12 hr light/dark cycle with *ad libitum* access to food and water. Working memory and body weights were monitored longitudinally, and more detailed phenotyping occurred at 6 and 14m. These time points were selected to obtain an adult phenotype (6m) and a middle-aged to aged time point (14m) that captured variation in disease symptoms before the mice starting exhibited severe health-related problems that confounded behavioral testing. All mouse experiments occurred at UTHSC and were conducted in accordance with the UTHSC Animal Care and Use Committee and the National Institutes of Health Guide for the Care and Use of Laboratory Animals.

## METHOD DETAILS

### Y-maze

The y-maze test of spontaneous alternation was performed as described previously (Oakley et al., 2006). The y-maze used for testing was made of clear acrylic with arms that were 2” wide x 12” long x 2” high. The maze was placed on a table in a dimly lit room and spatial cues were displayed on walls around the table. For all behavioral testing, mice were habituated to transport and to the testing room for three days prior to testing. Mice were placed in a randomized start arm and video tracking software was used to monitor arm entries (ANY-maze, Stoelting Co., IL, USA). An arm entry was called when the mouse’s entire body, including the two back feet, entered the arm. The sequence and total number of arms entered was recorded, and the percentage of successful alternations was calculated as follows: number of alternations/maximum possible alternations (total number of arms entered – 2) x 100. For each animal that was measured longitudinally (i.e. not harvested at the early 6mo time point), the age at which each animal became ‘impaired’, or performed below chance levels, was recorded and used as the animals “age at onset”. Strain averages for age at onset were determined and used for subsequent QTL mapping.

### Sensorimotor battery

At 6 and 14m, mice were subjected to a sensorimotor battery consisting of three tasks. First, mice were placed in the center of a 3-foot long narrow (0.5”) beam elevated 20.75” off a table surface and the time taken for the mouse to cross the narrow beam onto a safe platform on either side was measured. Second, mice were placed face-down on a wire mesh grid (holes were 1cm x 1cm) that was placed at a 45° angle. The time taken for a mouse to right itself (geotaxis) was recorded. A 3 minute maximum time limit was imposed for both the narrow beam and incline screen tests. If a mouse fell from the narrow beam, the maximum score of 180s was given. Third, grip strength was measured using a standard grip strength meter (Colbourn Instruments). Each of these three tasks were repeated in triplicate and the average score across three trials was used. For each mouse, a z-score based on the 6m population average was calculated for each task and the three z-scores were summed to derive a sensorimotor composite score.

### Elevated plus maze

At 6 and 14m, anxiety was evaluated using an elevated plus maze task. Mice were placed in the center of the maze and allowed to explore for 6 minutes. Video tracking software (ANY-maze, Stoelting Co.) was used to track the mouse and calculate the time spent in open versus closed arms of the maze as well as the number of arm entries into either open or closed arms, the total number of arm entries, and the total distance travelled in the maze.

### Contextual fear conditioning

Following 3 days of habituation to transport and to the testing room, mice were trained on a standard contextual fear conditioning paradigm as previously described (Neuner et al., 2015). Training consisted of a 180s baseline period followed by four mild foot shocks (1s, 0.9mA), separated by 115 ± 20s. A 40s interval following each foot shock was defined as the post-shock interval, and the percentage of time spent freezing during each of these intervals was measured using FreezeFrame software (Coulbourn Instruments, PA, USA). To calculate an acquisition curve for each strain as an index of learning, the slope across the average time spent freezing during post-shock intervals 1-4 was derived. Twenty-four hours later, hippocampus-dependent contextual fear memory (CFM) was tested by returning the mouse to the testing chamber for 10 min. The percentage of time spent freezing during the testing trial was measured using FreezeFrame software and used as an index of CFM. Pain sensitivity was evaluated in subset of mice by recording the length of activity burst following each shock. An average post-shock reactivity score was calculated by averaging the length of each activity bursts following the four training shocks.

### Enzyme-linked immunosorbent assay (ELISA)

Brains were removed at appropriate time points (6 or 14m) and immediately dissected, snap frozen, and stored at -80°C until use. Beta-amyloid 1-42 (Aβ42) levels were quantified from sections of cortex overlaying the hippocampus as previously described (Oakley et al., 2006). Briefly, tissue was homogenized in 1X PBS + 1% Triton-X 100 using the TissueLyser II system (Qiagen) and sonicated 2x10s on low power. Protein concentration was determined using a NanoDrop 2000 UV-Vis Spectrophotometer (ThermoScientific, USA). Brain homogenates (10 mg/ml) were extracted in a final concentration of 5M GuHCl overnight at 4°C. Samples were then diluted appropriately and run in duplicate on Aβ42-specific sandwich colorimetric ELISAs according to the manufacturer’s protocol (Cat# 298-92401, Wako Chemicals, Richmond, VA). Optical densities at 450 nm were read on a Biotek plate reader (BioTek, USA) and Aβ42 concentration was determined by comparison with Aβ42 standard curves. Only readings in the linear range of the standard curve were included in analysis. Duplicates were averaged to determine concentration of Aβ42 in each sample. Finally, Aβ42 concentrations were normalized to total protein concentration and are reported as nanograms of Aβ42 per milligrams of total protein.

### RNA sequencing

Snap frozen hippocampi from AD-BXD strains and non-carrier littermate controls at 6 and 14m were used for RNA sequencing. RNA was isolated on a Qiacube using the RNeasy mini kit (Qiagen) and treated with DNase to remove contaminating DNA. RNA quality was confirmed using a BioAnalyzer (Agilent Technologies). All samples had RNA Integrity Numbers (RIN values) > 8.0. Sequencing libraries were prepared from 1 μg RNA with the Truseq Stranded mRNA Sample Preparation Kit (Illumina Inc) following the manufacturer’s protocol. Final PCR-enriched fragments were validated on a 2200 Tapestation Instrument using the D1000 ScreenTape (Agilent Technologies) and quantified by qPCR using a Universal Library Quantification Kit (Kapa Biosystems) on the QuantStudio 6 Flex (ThermoFisher Scientific). Final library pools were sequenced by 75bp paired-end sequencing on a HiSeq2500 (Illumina Inc). Because both C57BL/6J and DBA/2J alleles segregate within our panel, the GBRS/EMASE pipeline developed by the Churchill group at The Jackson Laboratory was used in order to align reads to a diploid transcriptome (https://gbrs.readthedocs.io/en/latest/readme.html). An expectation maximization algorithm was used in order to align reads to the correct allele, allowing for the quantification of both total reads assigned to a gene and the number of reads assigned to either the *B* or *D* allele. For final by-strain analysis, samples belonging to the same strain/sex/age/genotype group were averaged. Pearson correlations to quantitative traits including age at onset, CFC acquisition, and CFM were performed using the cor function in R and corresponding p-values were generated using the corPvalueStudent function from the WGCNA package (Langfelder and Horvath, 2008). Differential expression analysis was conducted using the DESeq2 package (Love et al., 2014).

### Comparison of AD-BXD and human transcriptomes

In order to evaluate how well the AD-BXD transcript profile matches that of human AD, we utilized a dataset recently published by Hargis and Blalock (2017) comparing existing mouse models of AD to human AD. They identified a consensus AD signature consisting of 60 genes derived from the top 10% commonly upregulated and downregulated genes across three human AD datasets (**Table S2**). In order to see how the transcriptome from our AD-BXD panel compared to normal expression patterns, differential expression analyses comparing hippocampal gene expression from AD-BXD lines to non-carrier littermate controls was performed using DESeq2 (Love et al., 2014). Significantly differentially expressed genes were identified and a log2fold change was calculated for all genes that passed established filtering criteria (Love et al., 2014). The log_2_fold change (log_2_FC) for each of the 60 AD consensus genes was identified and used for comparison across human and mouse datasets obtained from Hargis et. al.. Differentially expressed genes were separated into those with a positive log2FC and those with a negative log2FC. These lists were then loaded into WebGestalt for overrepresentation enrichment analysis to identify pathways most different between AD- and Ntg-BXDs, and to compare these results to pathways affected in human AD.

### Genetic mapping

For males and females of each AD-BXD strain, averages were calculated for 1) age at onset, 2) contextual fear acquisition at 6 and 14m, 3) contextual fear memory at 6 and 14m, and 4) beta-amyloid 1-42 levels at 6 and 14m to use for genetic mapping using the Haley-Knott regression analysis in r/qtl (Broman et al., 2003). Genotypes for BXD strains (mm9) were downloaded from GeneNetwork.org and a table of genotypes for strains used here are available as **Table S7**. R/qtl’s scanone function using Haley-Knott regression and a single QTL model was used to identify regions of the genome containing variants responsible for regulating observed variation in quantitative traits (Broman et al., 2003). Sex and age were used as covariates where appropriate to allow for the possibility of different genetic contributions between sexes. The recommended 1.5 LOD confidence interval was identified and genes with sequence differences across C57BL6/J and DBA2/J, the two parental strains of our AD-BXD panel, in that interval were considered viable positional candidates. Sequences differences were identified using the Sanger Mouse Genomes Project mouse genomes variants querying site using the latest annotation (REL-1505). Ensembl Variant Effect Predictor (McLaren et al., 2016) was used to classify variants and insertions/deletions (indels) by predicted impact on gene product function.

### Hippocampal injections

Bilateral injections of AAV9 encoding either shRNA for *Trpc3* or a non-targeting control sequence were delivered into the hippocampus as previously described (Neuner et al., 2015). Briefly, mice were anesthetized using isoflurane gas and positioned in a stereotaxic frame. Guide cannula and a 28 gauge microinjector were used to deliver 1.0 uL of virus per hemisphere at a rate of 1.0 uL/min to the following coordinates: anterior-posterior -1.88 mm, medial-lateral ± 1.60, dorsal-ventral -1.46 mm. Mice were monitored and analgesic was administered every 12 hours for 72 hours post-surgery (Carprofen, 5-10 mg/kg).

### Western blot

Western blots were performed from whole hippocampal lysates from male 5XFAD-B6SJL mice as previously described (Neuner et al., 2015). Samples were removed from -80°C, placed in lysis buffer (0.32M sucrose, 3mM HEPES, pH 7.4 plus Roche miniComplete protease inhibitor cocktail) and homogenized using a rotor homogenizer (Kinematica, Switzerland). 15-20 μg of protein was loaded and separated on a 7.5% SDS-PAGE gel. Proteins were transferred using the Bio-Rad TurboTransfer system and blocked for 30 min at room temperature (RT). Primary antibodies for TRPC3 and GAPDH (Alamone Labs #ACC-016, Fitzgerald #10R-G109a respectively) were incubated overnight at 4°C and detected by anti-mouse and anti-rabbit fluorescent-conjugated antibodies. Visualization was performed using an Odyssey image scanner and blots were quantified using NIH ImageJ Software.

### Immunohistochemistry and image analysis

Immunohistochemistry for Aβ42 was performed in a subset of mice that had received AAV9 injections. Briefly, upon conclusion of testing, brains were dissected and one half was stored in 4% paraformaldehyde while the other half was subdissected and frozen in liquid nitrogen for future use. Fixed hemi-brains were embedded in agarose and 50 um coronal sections were cut on a vibrating microtome (Leica VT1000S). Sections were washed with PBS, incubated in 1% w/v sodium borohydride for 30 mins, washed again with PBS, and incubated in 5% normal donkey serum for 2 hours at room temperature. Primary antibody for Aβ42 (1:1000 dilution, Invitrogen, Cat #700254) was added and slices were incubated overnight at 4° C, washed with PBS, and secondary antibody (donkey anti-rabbit Alexa568, Invitrogen Cat# A10042) was added for 6 hours at RT. Slices were again washed with PBS and mounted with polyvinyl alcohol mounting medium with DABCO antifade (Fluka). Images from hippocampal CA1, dentate gyrus, and subiculum were captured at 10X using an epifluorescence microscope, the levels function in Adobe Photoshop was used identically across all images to increase visibility, and particle analysis in ImageJ was used to quantify the number of Aβ42-positive plaques.

### Analysis of human data

Data was obtained from the Religious Orders Study and Memory and Aging Project (ROS/MAP). ROS began in 1994 and involves older Catholic nuns, priests, and brothers recruited from across the US. MAP began in 1997 and involves older lay persons recruited from retirement communities, subsidized housing facilities, and social service agencies in the Chicago metropolitan area. Persons in both studies enrolled without dementia and agreed to annual clinical evaluations and organ donation at death (Bennett et al., 2012a; Bennett et al., 2012b). Immunohistochemistry was performed at autopsy and quantified using image analysis to calculate the percentage area of several brain regions occupied by Aβ (hippocampus, entorhinal, midfrontal, inferior temporal, angular, calcarine, ACC, and superior frontal cortices) (Bennett et al., 2012a; Bennett et al., 2012b). A final amyloid burden score was calculated by averaging the score across all eight regions. Clinical diagnosis was made at time of death based on consensus case conference (Schneider et al., 2007). We used data from clinical diagnosed AD patients who also exhibited amyloid pathology [defined as a Consortium to Establish a Registry for Alzheimer’s Disease (CERAD) neuritic plaque stage of “definite” or “probable” (Bennett et al., 2006; Mirra et al., 1991)] and correlated measures of amyloid burden with levels of *TRPC3* in the prefrontal cortex as measured by RNA-sequencing, collected as previously described (De Jager et al., 2014). Sex and age at death were adjusted for in our models. Gene expression data were calculated from prefrontal cortex tissue as part of the Accelerating Medicines Partnership AD project, and are publically available online (https://www.synapse.org/#!Synapse:syn2580853/wiki/). Other neuropathologies were calculated and described previously including lewy bodies (Schneider et al., 2012), PHF tau tangles (Bennett et al., 2012a; Bennett et al., 2012b), arteriolosclerosis (Buchman et al., 2011), atherosclerosis (Arvanitakis et al., 2017), cerebral amyloid angiopathy (Boyle et al., 2015), gross cerebral infarctions (Arvanitakis et al., 2011), and microinfarctions (Arvanitakis et al., 2011).

## QUANTIFICATION AND STATISTICAL ANALYSIS

All experiments were conducted with experimenters blind to strain background, genotype (5XFAD vs Ntg), or treatment (shRNA-Trpc3 vs shRNA-Ctrl) where appropriate. Similarly, data analysis was performed by experimenters blind to experimental condition. Statistical analysis was performed using SPSS software Version 23 (IBM) and R. Analyses included independent t-tests, univariate ANOVAs, Spearman or Pearson correlation, and permutation tests to determine statistical significance of QTL mapping results. Correction for multiple comparisons (Bonferroni correction) was also used where appropriate. Statistical tests on correlation coefficients were performed using Fisher’s r-to-z transformation as previously described. Most analyses were performed on a particular strain/sex/age group unless otherwise specified. Data values reported in both the main text and figure legends are given as mean ± standard error of the mean.

## DATA AND SOFTWARE AVAILABILITY

A table of BXD strains used in this study and their corresponding genotypes are available as Table S7. RNA-sequencing from the hippocampus of a subset of AD-BXD strains is available on Gene Expression Omnibus (GEO) under accession number GSE101144.

